# Single voxel autocorrelation uncovers gradients of temporal dynamics in the hippocampus and entorhinal cortex during rest and navigation

**DOI:** 10.1101/2021.07.28.454036

**Authors:** Nichole R. Bouffard, Ali Golestani, Iva K. Brunec, Buddhika Bellana, Jun Young Park, Morgan D. Barense, Morris Moscovitch

## Abstract

During navigation, information at multiple scales needs to be integrated. Single-unit recordings in rodents suggest that gradients of temporal dynamics in the hippocampus and entorhinal cortex support this integration. In humans, gradients of representation are observed, such that granularity of information represented increases along the long axis of the hippocampus. The neural underpinnings of this gradient in humans, however, are still unknown. Current research is limited by coarse fMRI analysis techniques that obscure the activity of individual voxels, preventing investigation of how moment-to-moment changes in brain signal are organized and how they are related to behavior. Here, we measured the signal stability of single voxels over time to uncover previously unappreciated gradients of temporal dynamics in the hippocampus and entorhinal cortex. Using our novel, single voxel autocorrelation technique, we show a medial-lateral hippocampal gradient, as well as a continuous autocorrelation gradient along the anterolateral-posteromedial entorhinal extent. Importantly, we show that autocorrelation in the anterior-medial hippocampus was modulated by navigational difficulty, providing the first evidence that changes in signal stability in single voxels are relevant for behavior. This work opens the door for future research on how temporal gradients within these structures support the integration of information for goal-directed behavior.

## Introduction

To enable efficient goal-directed behavior, information must be represented and integrated across multiple temporal and spatial scales. It has been proposed that neural signal gradients in the hippocampus and entorhinal cortex support such multi-scale representations in rodents, but evidence in humans is sparse and has methodological limitations. Previously, fMRI analysis techniques have uncovered local signal gradients in the human hippocampus (Brunec, Bellana, et al., 2018). These investigations, however, have been limited by analyzing patterns of activity across relatively coarse regions of interest, making it unclear how sustained versus rapidly changing signals are distributed throughout the hippocampus. Many of these analyses use predetermined anterior and posterior anatomical masks, which limit our ability to detect neural signal gradients in an unsupervised way, therefore preventing us from investigating gradients that exist along both anterior-posterior and medial-lateral axes of the hippocampus. Moreover, there have been no prior investigations of autocorrelation gradients in the entorhinal cortex, despite its key role in spatial and temporal representations during navigation. To address these limitations, we have developed a novel, data-driven analysis based on autocorrelation of single voxels in fMRI during rest and navigation. This technique allows us, for the first time, to track the signal stability of individual voxels and their spatial distribution in an unconstrained way along both the anterior-posterior and medial-lateral axes of the hippocampus and entorhinal cortex. Based on this single voxel analysis we uncover gradients of neural signal dynamics along these axes in both structures and relate them to behavior.

In rodents, place fields in the ventral hippocampus (homologous to the anterior hippocampus in humans) span larger areas, show a higher degree of overlap, and higher correlation in their firing across time, compared to the dorsal hippocampus (homologous to the posterior hippocampus in humans) (Hasselmo, 2008; Jung et al., 1994; Kjelstrup et al., 2008; Komorowski et al., 2013). A similar gradient of hippocampal organization is also observed in the human hippocampus. Tracking moment-to-moment similarity across patterns of voxels during virtual navigation, Brunec, Bellana, et al. (2018) found that signal similarity was significantly greater within the anterior hippocampus relative to the posterior hippocampus, indicating that, as in the rodent ventral hippocampus, the human anterior hippocampus demonstrates slower changing signals that are sustained across time and space. These results suggest that a relatively stable pattern of activity in the rodent and human hippocampus follows a scaled gradient, from faster changing signal in the posterior (dorsal) hippocampus to slower changing signal in the anterior (ventral) hippocampus. This gradient organization might underlie fine-to-coarse mnemonic representation, particularly when a different granularity of information needs to be maintained across time (Brunec & Momennejad, 2022; Robin & Moscovitch, 2017). In addition to the dorsal-ventral gradient of spatial representation observed in rodents, research suggests a difference in spatial selectivity along the proximodistal axis (homologous to medial-lateral in humans), specifically in CA1 (Igarashi et al., 2014), yet whether a similar medial-lateral distinction exists in the human hippocampus is still unclear (Hrybouski et al., 2019).

A key input structure to the hippocampus that has been implicated in integrating information over time during navigation is the entorhinal cortex. Prior research has found distinct functional differentiation between the anterolateral and posteromedial aspects of the entorhinal cortex (ERC), but there have been no prior investigations of neural signal gradients in the ERC. The lateral ERC in rodents, and the homologous anterolateral ERC in humans, supports within-object and object-location coding, as well as temporal information processing (Bellmund et al., 2019; Montchal et al., 2019; Olsen et al., 2017; Tsao et al., 2018; Yeung et al., 2017; 2019). In contrast, the posteromedial ERC in humans, has been primarily linked to scene processing (Berron et al., 2018; Maass et al., 2015; Navarro Schröder et al., 2015) and related to grid cell organization (Bellmund et al., 2016), consistent with evidence of grid cells in the medial ERC in rodents (Hafting et al., 2005). Given prior evidence of functional distinctions of the ERC into anterolateral and posteromedial regions, we developed a data-driven method to directly probe whether there exists a continuous neural signal gradient in both the anterior-posterior and medial-lateral axes of this structure.

Investigating these regions in tandem within the same analytic framework is also motivated by evidence of strong reciprocal connectivity between the hippocampus and ERC (Small et al., 2011; Maass et al., 2015; Dalton et al., 2019; Dalton et al., 2021).

To understand how a graded organization of signal dynamics in the hippocampus and ERC supports goal-directed behavior, we developed an analytic approach of temporal autocorrelation at the single voxel level, which we implemented during both rest and navigation. Temporal autocorrelation represents the degree of similarity between a signal and the temporally shifted, or lagged, version of the signal over successive time intervals (Figure 1A).

**Figure 1.**
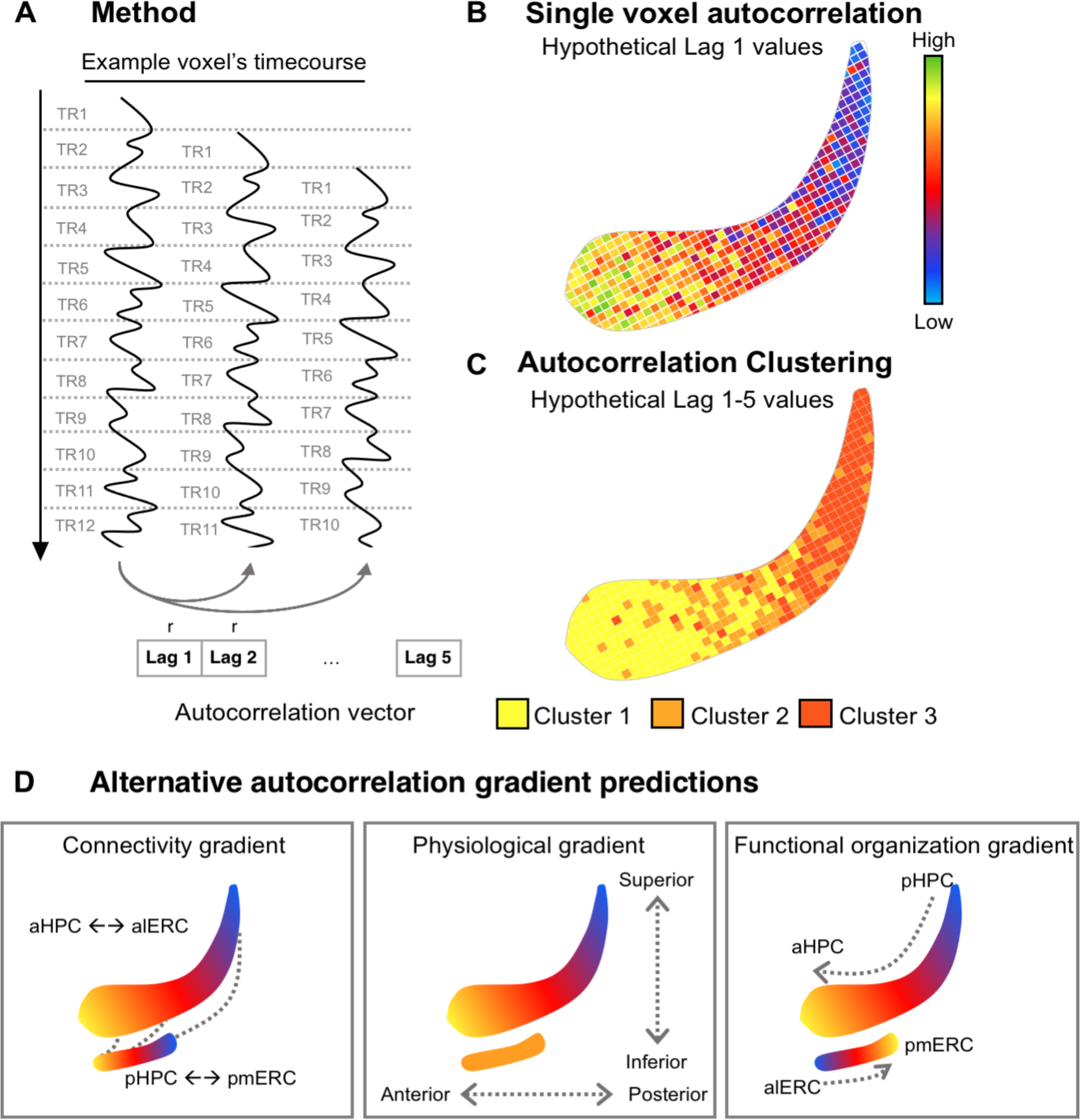
**A) Method.** For each voxel, the timecourse of activity was successively temporally shifted by 1 TR and correlated with itself. This was repeated for a total shift of 4 seconds (i.e., 5 lags for resting state data (Dataset 1) and 2 lags for navigation data (Dataset 2)). This resulted in a vector of single voxel autocorrelation values, with each value corresponding to a different lagged correlation. **B) Single voxel autocorrelation (hypothetical values).** The procedure was repeated for all voxels in an ROI. To examine the spatial distribution of the single voxel autocorrelation, we plot the group-level single voxel autocorrelation maps for each lag, averaged across runs and participants. **C) Autocorrelation clustering (hypothetical values).** The autocorrelation values for each lag were stored in a vector (single voxel autocorrelation vector). The voxels in the ROI were clustered based on the similarity (Euclidean distance) of single voxel autocorrelation vectors. Single voxel autocorrelation vectors were clustered according to their Euclidean distance (Blondel et al., 2008). Clustering was performed at the individual-level and at the group-level. **D) Three alternative autocorrelation gradient predictions.** One possibility is that autocorrelation gradients in the hippocampus and ERC would follow the pattern of reciprocal connectivity extending along the anterior-posterior extent. The second possibility is that autocorrelation gradients are driven by physiological or hemodynamic factors. If this were the case, autocorrelation values in the entorhinal cortex should be similar to those in the anterior portion of the hippocampus due to spatial proximity and similar anterior-posterior extent. The third possibility is that autocorrelation values in these structures result from intrinsic functional properties, which could give rise to differentiable patterns in hippocampus and ERC.

Conventionally, it is assumed that this autocorrelation in fMRI data originates from physical and physiological noise (Arbabshirani et al., 2014; Bollmann et al., 2018; Bullmore et al., 2001; James et al., 2019; Lenoski et al., 2008; Lund et al., 2006; Purdon & Weisskoff, 1998) or the hemodynamic response function (Arbabshirani et al., 2014; James et al., 2019; Rajapakse et al., 1998) and, therefore, has been considered irrelevant to brain function. Recently, however, Arbabshirani et al. (2019) found that autocorrelation reflects changes in cognitive state (task vs. rest) as well as changes in mental state (healthy control vs. schizophrenia), suggesting that the observed changes in the autocorrelation are also modulated by cognitive processes. Prior studies, however, have been limited and are unable to answer the question of how temporal autocorrelation is directly related to behavior. Examining the temporal autocorrelation of single voxels during an active navigation task, therefore, is important for understanding how a stable, highly correlated signal is relevant for behavior.

Investigating the fMRI signal at the single voxel level allows us to measure neural gradients with more precision than previous methods. While studies with fMRI in humans suggest that functional heterogeneity exists along the long axis of the hippocampus (Nadel et al., 2013; Poppenk et al., 2013; see Grady, 2020 for a review) and medial-lateral extent of the ERC (e.g., Hafting et al., 2005; Maass et al., 2015; Navarro Schröder et al., 2015), previous analysis techniques have been limited to investigations using predetermined anatomical masks, which obscures the contribution of individual voxels, making it unclear whether graded signals extend along multiple axes in these regions. Furthermore, examining the autocorrelation at the single voxel level allows for a finer-grained analysis that may be more sensitive to differences in navigational performance and can help us to determine how a scaled gradient of signal similarity might be employed to integrate representations across spatial scales during navigation.

We, therefore, combine our single voxel autocorrelation approach with an unconstrained clustering method to determine how temporal autocorrelation is distributed in multiple dimensions throughout the hippocampus and ERC.

Here we present evidence of a functional medial-lateral neural signal gradient in the hippocampus as well as a novel continuous gradient in the ERC. Using resting state fMRI data with high spatial and temporal resolution from the Human Connectome Project (HCP) (Dataset 1), we measured single voxel autocorrelation in the hippocampus and ERC. Using task-based fMRI (with lower spatial resolution) we measured the single voxel autocorrelation in the hippocampus while participants completed a real-world navigation task (Dataset 2). We measured the similarity of single voxels over time by correlating the timecourse of each voxel with temporally shifted versions of itself (Figure 1A).

We applied data-driven clustering to determine how temporal autocorrelation was spatially distributed throughout the hippocampus and ERC (Figure 1B-C). We reasoned that there were three plausible mechanisms that might drive distinct autocorrelation patterns (Figure 1D). The first possibility is that reciprocal connectivity between the hippocampus and ERC along the anterior-posterior axes drives the autocorrelation gradient. If this were the case, we would expect hippocampal and entorhinal gradients to follow an anterior-posterior organization. A second possibility is that autocorrelation gradients are primarily driven by physiological or hemodynamic factors that differ along the anterior-posterior and superior-inferior dimensions in the brain. In this scenario, autocorrelation values in the entorhinal cortex should be similar to those in the anterior hippocampus as it lies inferior to and extends slightly anterior to the hippocampus. The third – and most interesting – possibility is that the observed gradients reflect differences in the intrinsic organization and processing timescales of these regions, by way of their functional contributions to cognition. In this case, autocorrelation patterns between the hippocampus and ERC would be differentiable and related to behavior.

In support of the notion that autocorrelation gradients are driven by intrinsic functional properties of the different brain regions, we found clear evidence of differentiable autocorrelations patterns in hippocampus and ERC, and that hippocampal autocorrelation gradients were meaningfully related to behavior such that increases in navigation difficulty were associated with increases in autocorrelation in the anterior-medial hippocampus. These results are consistent with recent findings from resting-state MRI (deWael et al., 2018) and a meta-analytic framework (Genon et al., 2021) that suggest that there are two primary connectivity gradients in the hippocampus (one along the anterior-posterior axis and one along the medial-lateral axis). Importantly, we critically extend this work by showing that these gradients are related to behavior. Our single voxel autocorrelation approach yields consistent and precise gradients of single voxel autocorrelation in the hippocampus and ERC, providing a powerful new continuous and data-driven method that can illuminate how temporal dynamics in brain signals relate to complex cognition.

## Dataset 1: Resting state fMRI Materials and Methods *Participants*

We analyzed resting state fMRI data from 44 participants (14 male) from the Human Connectome Project (HCP) Retest dataset. This dataset consists of data from 44 participants who were scanned twice using the full HCP imaging protocol. All subject recruitment procedures and informed consent forms, including consent to share de-identified data, were approved by the Washington University Institutional Review Board (IRB) (Glasser et al. 2016). The present analysis of this dataset was approved by the University of Toronto research ethics board.

### Scanning parameters and preprocessing

Resting state data were collected using a multiband EPI pulse sequence (TR = 720 ms, TE = 33.1 ms, 72 slices with 2 mm thickness, FOV = 208 x 180 mm, voxel size = 2x2 mm, Flip angle = 52, Multiband factor = 8, Scan time = 14 minutes and 33 seconds). Each run was repeated twice, with a left-to-right and a right-to-left phase encoding direction. The presented results are generated from data with the left-to-right phase encoding direction.

Initial fMRI preprocessing steps already applied to the downloaded data included fieldmap correction, motion correction, brain extraction, registration to standard space, and intensity normalization (Glasser et al., 2013; Smith et al., 2013; Van Essen et al., 2013). The data were further preprocessed using the FIX tool in FSL (Griffanti et al., 2014; Salimi-Khorshidi et al., 2014), and noise components related to head motion and other artifacts were removed. To eliminate high frequency noise and artifacts, fMRI signals are low-pass filtered using MATLAB IIR Butterworth filter (designfilt function in Signal Processing Toolbox) with cutoff frequency of 0.1 Hz.

### Single voxel autocorrelation method

#### Computing single voxel autocorrelation

Bilateral hippocampal and entorhinal masks were generated using the Harvard-Oxford Atlas in FSL. For each voxel inside each of the regions of interest (ROIs), unbiased autocorrelation (as implemented in MATLAB xcorr function) was calculated. Specifically, the timecourse of a single voxel’s activity was correlated with itself shifted by a temporal lag, the length of 1 TR (Dataset 1 TR = 720 ms). We repeated this process, shifting the timecourse forward by 1 lag (720 ms) and correlating it with the original, non-shifted timecourse until a maximum temporal shift of 4 seconds was reached. We chose 4 seconds because it has been shown that the autocorrelation of the fMRI signal in the gray matter drops off after 4 seconds (i.e., it is not distinguishable from the autocorrelation of other noise) (Bollmann et al., 2018). For example, the non-shifted timecourse was correlated with lag 1 (length of 1 TR), lag2 (length of 2 TRs), etc. (Figure 1A). The autocorrelation (AC) computed for each lag was stored in a vector. The autocorrelation vector (single voxel autocorrelation vector) contained 5 values (one single voxel autocorrelation for each lag). This approach resulted in a single voxel autocorrelation vector for each voxel. All single voxel autocorrelation values were normalized by subtracting the mean and dividing by the standard deviation within each mask so that meaningful comparisons could be made between the two fMRI datasets (resting state and task). Single voxel autocorrelation maps were then averaged across the first and second runs from the 44 participants to generate an average overall map (e.g., Figure 1B).

#### Single voxel autocorrelation – Reliability Analysis

To verify that the observed single voxel autocorrelation pattern was not a measurement artifact (e.g., head motion, magnetic field inhomogeneity, physiological artifacts, etc.), we tested the reliability of the single voxel autocorrelation pattern within an individual. In our case, the single voxel autocorrelation pattern was deemed reliable if there was a high degree of agreement between the single voxel autocorrelation values generated from different runs from the same participant compared to runs from different participants. Reliability of the single voxel autocorrelation values was measured by calculating the Euclidean distance (ED) between the single voxel autocorrelation vectors for all pairs of run-wise datasets. 44 participants with 2 repeated sessions produced 44 intra-subject and 3784 inter-subject ED values. The lower the ED between two single voxel autocorrelation vectors, the higher the similarity between them. We expected to see more similar single voxel autocorrelation patterns between single voxel autocorrelation vectors generated from two runs of the same participant compared to two runs from different participants (lower intra-subject ED compared to inter-subject ED). The inter-subject and intra-subject ED are not completely independent from one another, therefore we used nonparametric permutation to test for significance. We randomly shuffled the intra-and inter-subject labels and pulled two samples of size 44 (intra-subject) and 3784 (inter-subject). We calculated the mean difference between the two samples and repeated this process 10,000 times, resulting in a histogram of mean differences under the null hypothesis (i.e., the difference between intra- and -inter-subject ED equal to zero). We compared the observed difference between intra- and inter-subject EDs with the null distribution and calculated nonparametric p-values. Permutation tests were conducted using a permutation testing package in Matlab (Laurens, 2021).

#### Computing single voxel autocorrelation clusters (Autocorrelation Clustering)

The Euclidean distance between the single voxel autocorrelation vectors of each voxel pair in each mask was calculated to create a distance matrix. The distance matrix was first normalized (i.e., divided by the maximum value) and then subtracted from 1 to generate a similarity matrix ranging from 0 to 1. This similarity matrix was used to generate hippocampal clusters by applying a Louvain clustering method using the modularity optimization algorithm proposed by (Blondel et al., 2008; Wickramaarachchi et al., 2014). Unlike the majority of the clustering methods, modularity optimization does not require to assign the number of clusters and estimates the optimum number of clusters from data. In addition to clustering at the level of each individual, group-level clustering was performed by averaging the similarity matrices of all participants (for a theoretical schematic, see Figure 1C).

#### Autocorrelation Clustering – Reliability Analysis

Reliability of the clustering was measured by calculating the overlap between the generated clusters using the Jaccard coefficient. The Jaccard coefficient of regions A and B is defined as:

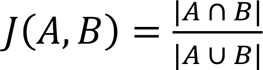

Where |*A* ∩ *B*| is the number of common voxels in both A and B (intersection) and |*A* ∪ *B*| is the number of voxels in A and B combined (union). Individual parcellations were then compared to the group-level parcellation to examine the consistency of parcellation. The Jaccard coefficient was calculated both intra-subject (overlap between clusters extracted from two runs from the *n*th subject) and inter-subject (overlap between the cluster from the *n*th subject and the same cluster estimated in all other subjects).

Assuming that the single voxel autocorrelation pattern is consistent across the two runs of the same participant, we expected there to be greater spatial overlap (higher Jaccard coefficient) among clusters within an individual compared to between different individuals. The Jaccard coefficients for clusters within participants are not completely independent from the Jaccard coefficients for clusters between participants, therefore we used nonparametric permutation to test for significance. For each cluster, we randomly shuffled the intra-and inter-subject labels and pulled two samples of size 44 (intra-subject) and 3784 (inter-subject). We calculated the mean difference between the two samples and repeated this process 10,000 times, resulting in a histogram of the mean differences under the null hypothesis (i.e., the difference between intra- and - inter-subject Jaccard coefficient equal to zero). We compared the observed difference between intra- and inter-subject Jaccard coefficients with the null distribution and calculated nonparametric P values.

#### Temporal SNR Control Analysis

Temporal SNR is a measure of noise that may interfere with the estimation of the autocorrelation of the fMRI signal and varies in intensity throughout the brain (Hutton et al., 2011). To rule out the possibility that the hippocampal autocorrelation clusters are being driven by noise, we calculated the temporal SNR for each cluster and tested whether the hippocampal autocorrelation could be explained by temporal SNR. We computed the temporal SNR for each voxel by taking the average of the voxel’s timecourse and dividing it by the standard deviation. The fMRI data used to calculate the tSNR did not have temporal filtering applied. We averaged the temporal SNR across all the voxels in each cluster to get the average temporal SNR per cluster. We then conducted linear mixed effects models predicting average autocorrelation of each cluster (lag 1) with Hemisphere (left,right), Cluster(1,2,3), and average temporal SNR as fixed effects and participant as the random intercept in the random effects term.

## Results Hippocampus

### Spatial distribution of single voxel autocorrelation

Our analysis correlated the timecourse of activity of each voxel in the hippocampus with activity in that same voxel shifted by a temporal lag of 1 TR, repeating this process until a maximum temporal shift of 4 seconds (5 lags) was reached (Figure 1A) and generated maps of single voxel autocorrelation values for each lag (for a theoretical schematic, see Figure 1B). We found that single voxel autocorrelation maps at the group level (lags 1-5) showed a notable difference in the distribution of single voxel autocorrelation values along hippocampal axes (Figure 2A). More specifically, voxels with higher single voxel autocorrelation were mainly in the anterior-medial region whereas voxels with lower single voxel autocorrelation were mainly in the posterior-lateral region (in both left and right hippocampus). As shown in Figure 2A, although the overall autocorrelation decreased as the lag increased, the overall pattern of autocorrelation gradients was similar for lags 1-5.

**Figure 2.**
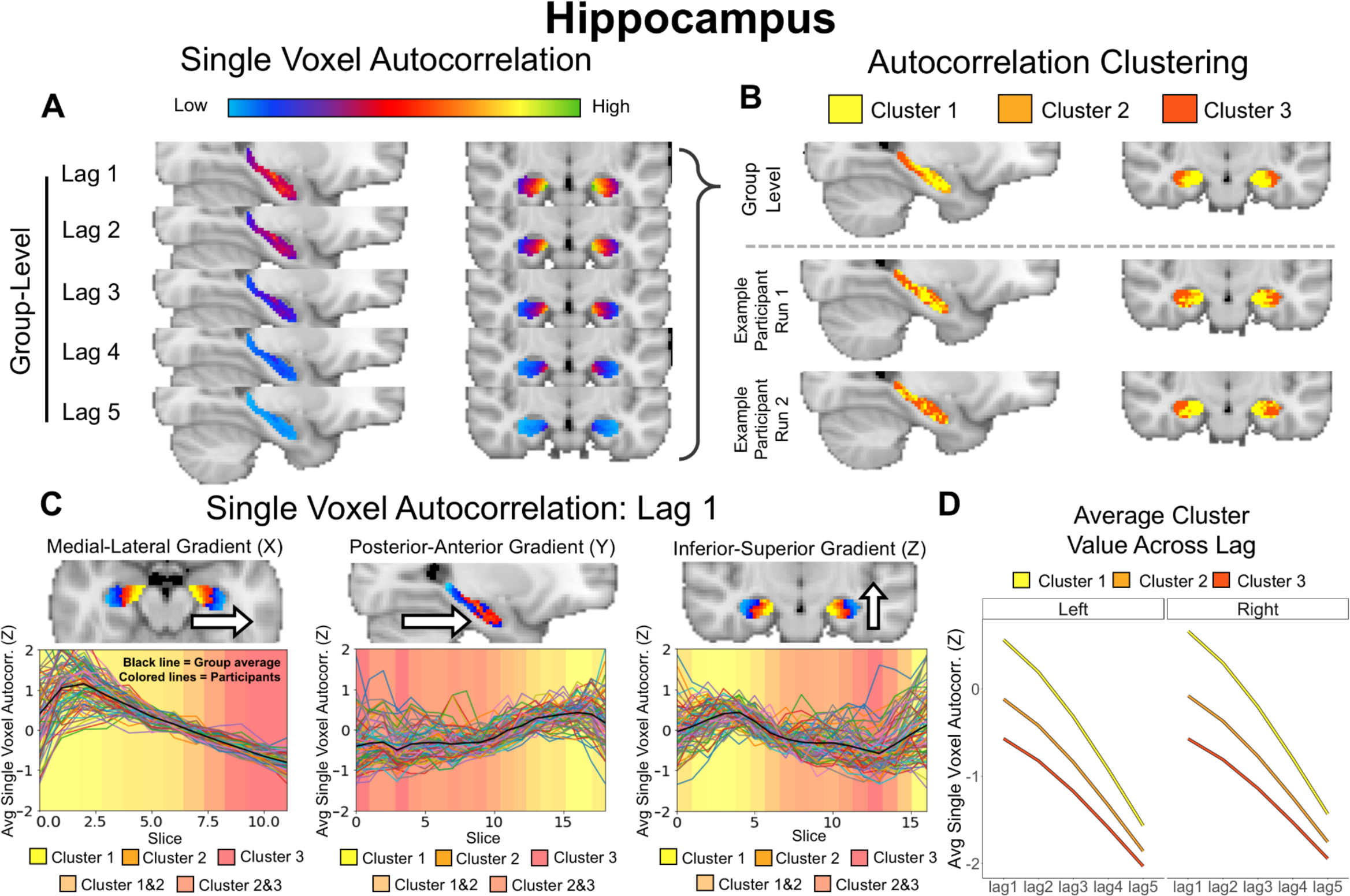
**Hippocampus. Single Voxel Autocorrelation. A)** Group-level single voxel autocorrelation maps averaged across all runs for all participants. **Autocorrelation Clustering. B)** Group-level clusters (top) and run-level cluster maps for two runs from an example participant (bottom). Three distinct clusters were found at both the group and the individual run-level. Cluster 1 was located in the anterior-medial hippocampus, Cluster 3 was located in the posterior-lateral hippocampus, and Cluster 2 was located between Cluster 1 and 3. **C) Single Voxel Autocorrelation: Lag 1.** Single voxel autocorrelation (lag 1) averaged per slice and projected into three axes (X, Y, and Z) to visualize changes in medial-lateral, anterior-posterior, and inferior-superior directions (plots depict left hemisphere; right hemisphere looked similar). The average cluster assignment of voxels on each slice is presented as the background color to show the gradation in values along the three axes. **D) Average Cluster Value Across Lag.**

### Single voxel autocorrelation – Reliability results

We next tested the reliability of these results. Here we defined a reliable result as one in which single voxel autocorrelation vectors generated from two runs of the *same* participant were more similar than two runs from *different* participants (lower intra-subject Euclidean distances compared to inter-subject Euclidean distances). Nonparametric permutation tests comparing intra-subject and inter-subject Euclidean distance revealed reliable results in both the left (intra-subject: 7.43 ± 2.07, inter-subject: 9.38 ± 2.22, P < 0.0001) and right hippocampus (intra-subject: 7.22 ± 1.98, inter-subject: 9.02 ± 2.02, P < 0.0001) (Supplemental materials, S1 panel A). These high intra-subject similarity values suggest that the single voxel autocorrelation pattern is an intrinsic feature of the brain, likely originating from neuronal sources, rather than noise or imaging artifacts.

### Autocorrelation Clustering

Data-driven clustering of the single voxel autocorrelation vectors revealed three distinct clusters in both the left and right hippocampus (Figure 2B); notably, past work that segmented the hippocampus into two ROIs (anterior and posterior) a priori would not have been able to detect the presence of this third cluster. Consistently across all 5 lags we found that Cluster 1 had the highest single voxel autocorrelation values and was located in the anterior-medial hippocampus (Figure 2D). Cluster 3 had the lowest single voxel autocorrelation values and was located in the posterior-lateral part of the hippocampus. Cluster 2 had intermediate single voxel autocorrelation values and was located between Clusters 1 and 3. These three clusters were also reliably observed at the individual level (cluster maps from two runs of an example participant are shown in Figure 2B).

In summary, clustering revealed a high-to-low single voxel autocorrelation gradient along the anterior-posterior axis, consistent with what has been previously found in the literature (Brunec, Bellana, et al., 2018; Raut et al., 2020). In addition, we found differences along the medial-lateral axis, as well as a prominent anterior-medial cluster of high single voxel autocorrelation that could be distinguished from a posterior-lateral cluster of low single voxel autocorrelation. While previous methods using predetermined anterior/posterior ROI masks might have missed this medial-lateral distinction, our data-driven method provides evidence that an autocorrelation gradient exists along multiple spatial dimensions.

Average group-level single voxel autocorrelation values for each cluster at each lag. Cluster 1 was associated with the highest single voxel autocorrelation values, Cluster 2 with intermediate values, and Cluster 3 with the lowest. This was consistent across all 5 lags.

### Autocorrelation Clustering – Reliability results

The reliability of single voxel autocorrelation clustering was evaluated by measuring spatial overlap between clusters, calculated by the Jaccard coefficient (Supplemental materials, S1 panel B). Here we defined a reliable result as one in which the spatial distribution of autocorrelation clusters was consistent across the two runs of the same participant, indicated by greater overlap (higher Jaccard coefficient) among clusters *within* an individual compared to *between* different individuals. Using nonparametric permutation, we found high reliability for clusters in the bilateral hippocampus, specifically Cluster 1 (Left: P < 0.001; Right: P < 0.001) and Cluster 3 (Left: P < 0.001; Right: P < 0.001). These findings of high intra- and inter-subject overlap suggest that Clusters 1 and 3 were highly reliable, within individuals. Cluster 2, however, had significantly lower overlap (Left: P = 0.06; Right: P = 0.007), suggesting more variability within individuals.

#### Temporal SNR Control Analysis

To rule out the possibility that the hippocampal autocorrelation clusters are being driven by noise, we calculated the temporal SNR for each cluster and tested whether the hippocampal autocorrelation could be explained by temporal SNR. We ran a linear mixed effects model with Hemisphere (left, right), Cluster (1,2,3) and temporal SNR as fixed effects predictors and average autocorrelation of each cluster (lag 1) as the predicted variable. We found a significant effect of Cluster (F(2,461.84) = 6619.86, p < .001), suggesting that even when temporal SNR is included in the model, clusters significantly explained the variance of the autocorrelation. We did not find a significant effect of temporal SNR. Additionally, we ran another model without Cluster as a fixed effect predictor and compared the two models (the one with and without Cluster) using a Chi square test. We found that the model including Cluster was significant X^2^ (8, N = 19) = 1422.9, p < .001, indicating that the model including Cluster was a better fit of the data compared to the model that only had hemisphere and temporal SNR alone. Together these results suggest that temporal SNR alone cannot explain the hippocampal autocorrelation clusters.

### Gradients of single voxel autocorrelation (lag 1)

The single voxel autocorrelation and autocorrelation clustering results presented above both suggest the presence of an autocorrelation gradient along two main axes: the anterior-posterior axis and the medial-lateral axis. To examine these individual gradients more precisely, we plotted the single voxel autocorrelation across hippocampal slices along the X (medial-lateral), Y (posterior-anterior), and Z (inferior-superior) axes. We observed consistent gradients in every participant. Specifically, single voxel autocorrelation gradually decreased in the medial-to lateral direction and increased in the posterior-to-anterior direction (Figure 2C; we focused on lag 1, but a similar pattern was revealed across all lags, as shown in Figure 2A). A rough gradient of high-to-low autocorrelation was also observed in the inferior-superior axis, which is due to the angle of the hippocampus (i.e., the anterior hippocampus is located more inferiorly relative to the posterior hippocampus). When we investigated the spatial distribution of the three clusters (projected on the background of the plots in Figure 2C), we observed a gradient of cluster assignment that complemented the single voxel autocorrelation gradients. Specifically, high-to-low single voxel autocorrelation gradients were also associated with a cluster gradient from Cluster 1 to Cluster 3.

### Entorhinal cortex

#### Spatial distribution of single voxel autocorrelation

We repeated the analyses above in the ERC. To illustrate the distribution of autocorrelation values of individual voxels throughout the ERC, we plotted the group-level single voxel autocorrelation maps for lags 1-5 (Figure 3A). The maps illustrate a difference in single voxel autocorrelation throughout the ERC. Specifically, voxels with higher single voxel autocorrelation were mainly in the posterior-medial region whereas voxels with lower single voxel autocorrelation were mainly in the anterior-lateral region (in both left and right ERC).

**Figure 3.**
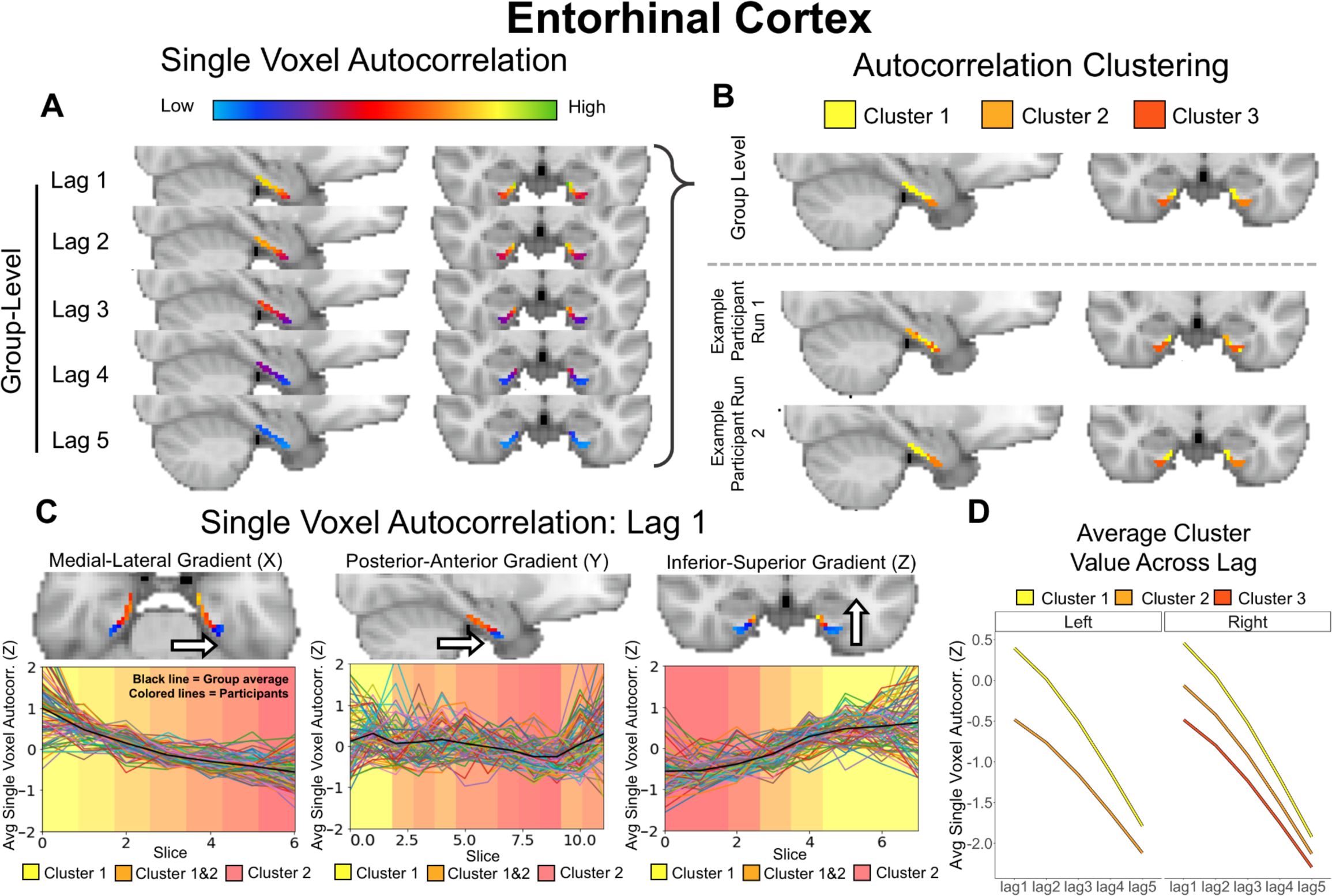
**Entorhinal cortex. Single Voxel Autocorrelation. A)** Group-level single voxel autocorrelation maps averaged across all runs for all participants. **Autocorrelation Clustering. B)** Group-level clusters (top) and run-level cluster maps for two runs from an example participant (bottom). Two distinct clusters were found in the left hemisphere and three in the right hemisphere. In the left hemisphere, Cluster 1 was located in the posterior-medial ERC and Cluster 2 was in the anterior-lateral ERC. In the right hemisphere Cluster 1 was located in the posterior-medial ERC, Cluster 3 was located in the anterior-lateral ERC, and Cluster 2 was located between Cluster 1 and 3. **C) Single Voxel Autocorrelation: Lag 1.** Single voxel autocorrelation projected below onto three axes (X, Y, and Z) to visualize changes in medial-lateral, anterior-posterior, and inferior- superior directions (for the left hemisphere; right hemisphere looked similar). The average cluster assignment of voxels on each slice is presented as the background color to show the gradation in values along the three axes (Note the gradation depicts only Cluster 1 and 2 as there were only two significant clusters found in the left hemisphere). **D) Average Cluster Value Across Lag.** Average group-level single voxel autocorrelation values for each cluster at each lag. In the left hemisphere, Cluster 1 was associated with the highest single voxel autocorrelation values and Cluster 2 with low autocorrelation values. In the right hemisphere, Cluster 1 was associated with the highest single voxel autocorrelation, Cluster 2 with intermediate values, and Cluster 3 with the lowest. This was consistent across all 5 lags.

#### Single voxel autocorrelation – Reliability results

Nonparametric permutation tests comparing intra-subject and inter-subject Euclidean distance revealed reliable results in both the left (intra-subject: 11.43 ± 3.64, inter-subject: 12.87 ± 2.83, P < 0.001 and right ERC (intra-subject: 10.41 ± 2.63, inter-subject: 13.19 ± 2.53, P < 0.001; see supplemental materials, S1 panel C). This analysis demonstrates the reliability of the single voxel autocorrelation and suggests that single voxel autocorrelation patterns between vectors generated from two runs of the *same* participant were more similar than two runs from *different* participants (lower intra-subject Euclidean distances compared to inter-subject Euclidean distances).

### Autocorrelation Clustering

The group-level clustering analysis on the voxels within the ERC revealed two distinct clusters in the left hemisphere and three clusters in the right (Figure 3B). For comparison, cluster maps from two runs of an example participant are shown in Figure 3B. Cluster 1 was located in the posteromedial ERC and had the highest single voxel autocorrelation values in both left and right hemispheres.

Cluster 2 was observed in the left hemisphere and was located towards the anterior-lateral ERC with low single voxel autocorrelation values. In the right hemisphere it was an intermediate cluster. Cluster 3 was only observed consistently in the right hemisphere and was located in the anterior-lateral ERC with the lowest single voxel autocorrelation values. We computed the group-level single voxel autocorrelation for each cluster and plotted it across all 5 lags (Figure 3D). Across all 5 lags, Cluster 1 consistently had the highest single voxel autocorrelation values, followed by Cluster 2 and Cluster 3.

### Autocorrelation Clustering – Reliability results

The reliability measure for ERC clusters was calculated by the Jaccard coefficient (Supplemental materials, S1 panel D). Nonparametric permutation tests comparing intra-subject and inter-subject cluster overlap revealed reliable results in the left and right hemisphere. In the left hemisphere, the Cluster 1 (P < 0.001) and Cluster 2 (P < 0.001) were reliable. In the right hemisphere, Cluster 1 (P < 0.001) and Cluster 3 (P < 0.001) were reliable. This suggests that these clusters were highly reliable within individuals. In the right hemisphere, Cluster 2 had very small Jaccard values, suggesting less reliability within individuals (Right: P = 0.53).

#### Temporal SNR Control Analysis

We calculated the temporal SNR for each cluster and tested whether the entorhinal autocorrelation clusters could be explained by temporal SNR. We ran a linear mixed effects model with Hemisphere (left, right), Cluster(1,2,3) and temporal SNR as fixed effects predictors and average autocorrelation of each cluster (lag 1) as the predicted variable. We found a significant effect of Cluster (F(2,396.42) = 1237.65, p < .001), suggesting that even when temporal SNR is included in the model, clusters significantly explained the variance in the autocorrelation. We did not find a significant effect of temporal SNR. Additionally we ran another model without Cluster as a fixed effect predictor and compared the two models (the one with and without Cluster) using a Chi square test. We found that the model including Cluster was significant X^2^ (8, N = 19) = 781.62, p < .001, indicating that the model including Cluster was a better fit of the data compared to the model that only had hemisphere and temporal SNR alone. Together these results suggest that temporal SNR alone cannot explain the entorhinal autocorrelation clusters.

### Gradients of single voxel autocorrelation (lag 1)

Single voxel autocorrelation values for lag 1 were projected onto X (medial-lateral), Y (posterior-anterior), and Z (inferior-superior) axes. As shown in Figure 3C, in every participant, single voxel autocorrelation values gradually decreased in the medial-to-lateral direction and the posterior-to-anterior direction (Figure 3C). We found a gradient of low-to-high autocorrelation along the inferior-superior axis, which is due to the fact that the posterior region of the ERC is more superior than its anterior region. We observed a gradient of cluster assignment that complemented the single voxel autocorrelation gradients, where high-to-low gradients were also associated with a cluster gradient from Cluster1 to Cluster 2.

### Dataset 2: Navigation *fMRI*

We next aimed to replicate the observed effects in task fMRI and relate changes in single voxel autocorrelation to behavior. Specifically, we were interested in how single voxel autocorrelation throughout the hippocampal long axis might be modulated by differences in difficulty during a temporally-extended navigation task. Therefore, we performed our single voxel autocorrelation analyses on a task fMRI dataset acquired while participants navigated in a familiar virtual reality environment (previously described in Brunec, Bellana et al.,

2018). Due to the lower spatial resolution in this dataset, we were not able to examine the ERC and, thus, these analyses focused only on the hippocampus.

## Material and Methods

### Participants

Nineteen participants (9 males; mean age 22.58 years, range 19-30 years) were scanned while navigating Google Street View routes around the city of Toronto. All subject recruitment procedures and informed consent was approved by the University of Toronto research ethics board.

### Paradigm

Participants met with the experimenter ahead of time and built routes that were either highly familiar or less familiar to them (e.g., frequently traveled or not). Participants then returned to the lab for their second session and were scanned while they navigated four different types of routes. 1) Familiar: participants started at a familiar landmark and navigated to a familiar goal destination via a familiar route, 2) Mirrored: participants started at a familiar landmark and traveled to a familiar destination via a familiar route, but the images of the route were mirrored (left-right reversed), 3) Unfamiliar: participants started at a familiar location, navigated to a familiar destination, but they were instructed to take an unfamiliar route between the two, and 4) GPS: participants started at an unfamiliar location in an unfamiliar part of town and followed the directions displayed by an arrow on the screen to the goal destination.

Participants completed four unique routes in each condition, sixteen routes in total (1 route = 1 scanned run). At the end of each route, participants rated the difficulty of the route on a scale from 1 (difficult) to 9 (easy).

### Scanning parameters and preprocessing

Participants were scanned with a 3T Siemens MRI scanner at Baycrest’s Rotman Research Institute. A high-resolution 3D MPRAGE T1-weighted pulse sequence image (160 axial slices, 1 mm thick, FOV = 256 mm) was first obtained to register functional maps against brain anatomy. Functional imaging was performed to measure brain activation by means of the blood oxygenation level dependent (BOLD) effect. Functional T2*-weighted images were acquired using echo-planar imaging (30 axial slices, 5 mm thick, TR = 2000 ms, TE = 30 ms, flip angle = 70 degrees, FOV = 200 mm). The native EPI resolution was 64 x 64 with a voxel size of 3.5mm x 3.5mm x 5.0mm. Images were first corrected for head motion using the Analysis of Functional NeuroImages (AFNI; Cox, 1996). All subsequent analysis steps were conducted using the statistical parametric mapping software SPM12.

Preprocessing involved slice timing correction, spatial realignment and co-registration, with a resampled voxel size of 3mm isotropic, with no spatial smoothing. As all of our analyses rely on covariance, we additionally regressed out the mean time-courses from participant-specific white matter, and cerebrospinal fluid masks, alongside estimates of the 6 rigid body motion parameters from each EPI run. To further correct for the effects of motion which may persist despite standard processing (Power et al., 2012), an additional motion scrubbing procedure was added to the end of our preprocessing pipeline.

Using a conservative multivariate technique, time points that were outliers in both the six rigid-body motion parameter estimates and BOLD signal were removed, and outlying BOLD signal was replaced by interpolating across neighboring data points. Motion scrubbing further minimizes any effects of motion-induced spikes on the BOLD signal, over and beyond standard motion regression, without leaving sharp discontinuities due to the removal of outlier volumes (for details, see Campbell et al., 2013). To enable comparisons at the group-level, the final step of the preprocessing involved warping participants’ functional data to the MNI-space template.

### Single voxel autocorrelation method

#### Computing single voxel autocorrelation

To compute the single voxel autocorrelation, we completed the same procedure outlined in Dataset 1. We used the same bilateral hippocampal masks to extract the HPC voxels in Dataset 2. For each voxel, the single voxel autocorrelation was calculated by repeatedly shifting temporal lags (length of 1 TR) until a maximum lag of 4 seconds was reached. In Dataset 2 the TR was 2000 ms; therefore, single voxel autocorrelation for 2 lags (or 2 TRs) was calculated, resulting in a maximum lag of 4 seconds. As outlined in the procedure above, single voxel autocorrelation values were normalized by subtracting the mean and dividing by the standard deviation. Single voxel autocorrelation was calculated for all four runs of each navigational condition (Familiar, Unfamiliar, Mirrored, GPS). The single voxel autocorrelation was averaged across the four scanned runs (unique routes), resulting in four different maps (one for each navigational condition). Single voxel autocorrelation maps were then averaged across the 19 participants to generate an average group map for each navigation condition.

Participants completed 16 navigation runs (four in each condition) at their own pace. Because the conditions varied in difficulty, the average number of TRs differed across conditions and participants. Every route was 2-10 km long and the average run (route) length was 137.6 TRs (2 s TRs). The average number of TRs was lowest in the GPS condition (M = 92.13, SD = 17.44), followed by the Familiar condition (M = 136.45, SD = 39.18), the Mirrored condition (M = 155.73, SD = 36.84), and the Unfamiliar condition (M = 158.78, SD = 32.13). In order to compare single voxel autocorrelation across scanned runs with a similar number of TRs/lengths, we chose to filter out any runs that were unusually short (that the participant either didn’t complete or completed very quickly). We excluded runs that were less than 88 TRs long. This resulted in an average of 13.36 runs (SD = 1.21) per participant. The GPS runs were disproportionately shorter than the other conditions, resulting in more GPS runs excluded than other conditions. For two participants, all four control runs were less than 88 TRs long, and therefore all of their control runs were excluded. The average number of routes included in the following analyses per participant are as follows: Mirrored (M=3.89, SD=0.31), Unfamiliar (M=3.84, SD=0.50), Familiar (M=3.68, SD=0.47), GPS (M=1.95, SD=1.22).

#### Computing autocorrelation clusters (Autocorrelation Clustering)

We repeated the single voxel autocorrelation clustering procedure described above in Dataset 1 to determine clusters of single voxel autocorrelation within each navigational condition.

### Relating single voxel autocorrelation to navigation condition

#### Comparing average autocorrelation of clusters across navigation condition

To investigate how the spatial distribution of single voxel autocorrelation is related to navigation difficulty, we compared the average single voxel autocorrelation (lag 1) of each cluster across the four different conditions: GPS, Familiar, Unfamiliar, and Mirrored. For each participant, we averaged the single voxel autocorrelation (lag 1) across all voxels in each cluster and Z transformed the values. To test whether there was a significant difference between single voxel autocorrelation during different navigation conditions, we modeled the data using linear mixed effects models with fixed effects based on experimental hypothesis and random effects structures to account for the effects of individual participants. We included Hemisphere, Cluster, and Navigation condition as fixed effects and participant as the random intercept in the random effects term. We began each model with a random effects structure that was the maximal justified by the design (including random slopes) (Barr et al., 2013). We systematically pruned the random effects structure until the model converged while avoiding a singular solution (i.e., overfitting) (Singmann et al., 2019). Models that included random slopes either failed to converge or reached a singular solution, therefore the models reported only include random intercepts for random effects term.This analysis was conducted in R (R Core Team, 2019) using the afex (Singmann et al., 2020) and the tidyverse packages (Wickham, 2017). We used the Tukey method to correct p-values for multiple comparisons in post-hoc tests.

#### Temporal SNR Control Analysis

We conducted the control analysis described in Dataset 1 to make sure that the difference in hippocampal autocorrelation across navigation conditions that we observed could not solely be explained by temporal SNR. We ran a linear mixed effects model on the average autocorrelation per cluster (lag 1) with Hemisphere (left, right), Cluster(1,2,3), Navigation condition (GPS, Familiar, Unfamiliar, Mirrored), and temporal SNR per cluster as fixed effects predictors.

## Results

### Hippocampus

#### Spatial distribution of single voxel autocorrelation

We observed a difference in single voxel autocorrelation along the anterior-posterior and medial-lateral hippocampal axes, where voxels with higher single voxel autocorrelation were found in the anterior-medial hippocampus and voxels with lower single voxel autocorrelation were found in the posterior-lateral hippocampus. Figure 4A shows the group-level single voxel autocorrelation maps for the four navigation conditions (as single voxel autocorrelation maps for lags 1- 2 were similar, only lag 1 is depicted in Figure 4A). The spatial distribution of single voxel autocorrelation was similar across navigation conditions and was also similar to the findings from Dataset 1. In the next section, we investigate the differences between conditions in more depth.

**Figure 4.**
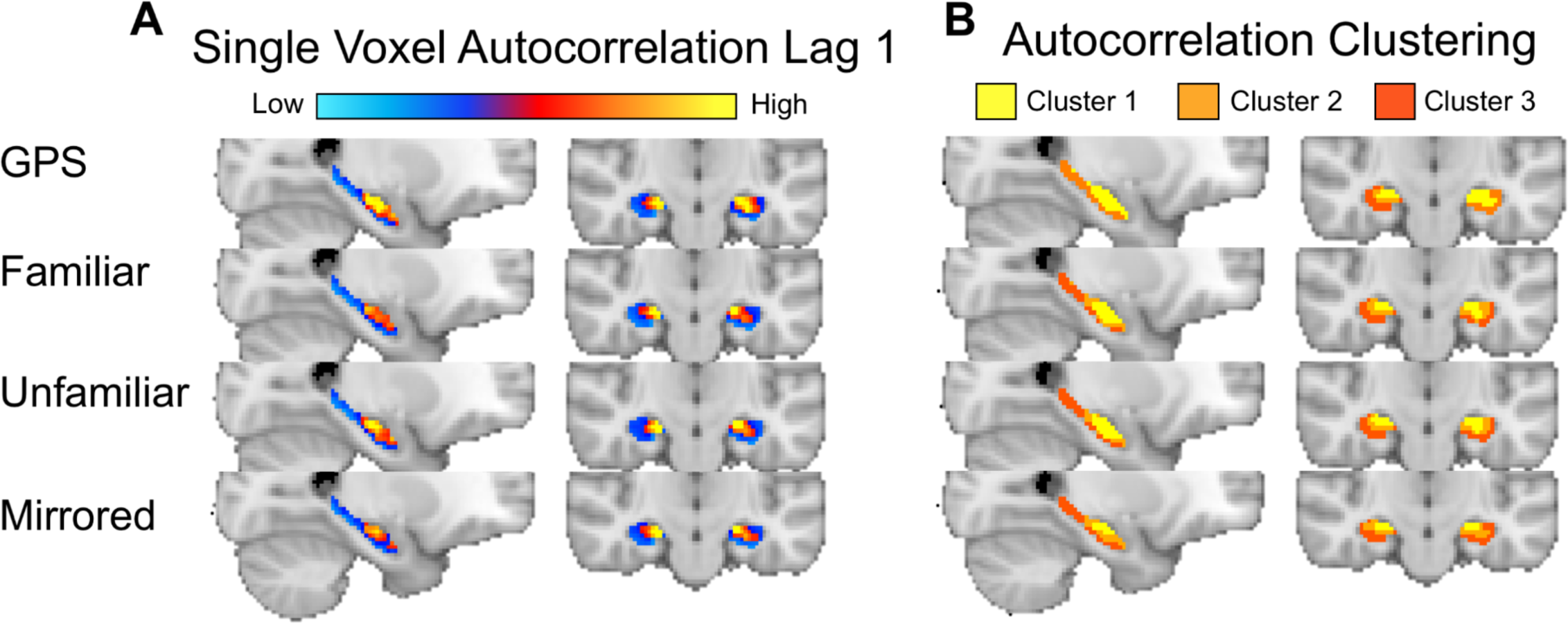
**A) Single Voxel Autocorrelation: Lag 1.** Single voxel autocorrelation values at lag 1 for every voxel in the hippocampus during spatial navigation. These values are averaged across run and participant for each of the GPS, Familiar, Unfamiliar and Mirrored conditions. A gradient from high to low autocorrelation is observed in the anterior-posterior and medial-lateral axes, across all navigation conditions. **B) Autocorrelation Clustering.** Cluster maps averaged across run and participant for each route type. High single voxel autocorrelation voxels cluster in the anterior-medial hippocampus and low single voxel autocorrelation voxels cluster in the posterior-lateral hippocampus.

#### Autocorrelation Clustering

As with Dataset 1, autocorrelation clustering revealed three distinct clusters in the left and right hemispheres for the Familiar, Unfamiliar and Mirrored conditions (Figure 4B). For Familiar, Unfamiliar, and Mirrored conditions, Cluster 1 was located in the anterior-medial HPC and had the highest single voxel autocorrelation. Cluster 3 was located in the posterior-lateral hippocampus and had the lowest single voxel autocorrelation. Cluster 2 was located between Cluster 1 and 3 and had intermediate single voxel autocorrelation. The GPS condition had three clusters in the right hemisphere and only two in the left.

#### Relating single voxel autocorrelation to navigation condition

Subjective difficulty ratings collected after each route (1 = difficult, 9 = easy) suggested that across the navigation conditions, navigational difficulty increased. Participants rated the GPS routes as the easiest (M = 7.2, SD = 1.46), followed by the Familiar condition (M = 6.98, SD = 2.05), Unfamiliar condition (M = 4.35, SD = 2.66), and the Mirrored condition, which was subjectively the most difficult (M = 3.97, SD = 2.42).

As navigation becomes more difficult, it is beneficial to integrate or maintain information over time, which may be reflected in changes in single voxel autocorrelation. Specifically, more stable neural dynamics might enable individuals to maintain information as one moves towards a goal. This prediction leads to two possibilities. In the first, as navigational difficulty increases we might observe a uniform change in single voxel autocorrelation across all voxels in the hippocampus. A second possibility is that as difficulty increases, voxels that tend to exhibit high autocorrelation during rest would differentially increase their autocorrelation relative to voxels that tend to exhibit low autocorrelation. To investigate these possibilities, we calculated the average single voxel autocorrelation (lag1) of each of the three clusters for each navigation condition. If navigational difficulty leads to a uniform increase in autocorrelation, we would observe no interaction between cluster and navigation condition. However, if navigational difficulty disproportionately affects the regions of the hippocampus that show high autocorrelation during rest (Figure 2), then more difficult routes would elicit a larger change in autocorrelation in the anterior-medial cluster (Cluster 1) compared to the more intermediate or posterior-lateral clusters (Clusters 2 and 3).

#### Comparing average autocorrelation of clusters across navigation condition

We compared the average single voxel autocorrelation of each cluster across the four route conditions: GPS, Familiar, Unfamiliar, and Mirrored. In both the left and right hemisphere, across all four navigation conditions, Cluster 1 had the highest autocorrelation (*Left:* M = 1.00, SD = 1.27; *Right:* M = 0.77, SD = 1.44) compared to Cluster 2 (*Left:* M = -0.11, SD = 0.64; *Right:* M = -0.21, SD = 0.51) and Cluster 3 (*Left:* M = -0.55, SD = 0.11; *Right:* M = -0.55, SD = 0.13), which is consistent with our findings from the autocorrelation clustering in Dataset 1.

To test whether there was a significant difference between single voxel autocorrelation during different navigation conditions, we ran a linear mixed effects model on the autocorrelation of the clusters. We included Hemisphere, Cluster, and Navigation condition (GPS, Familiar, Unfamiliar, and Mirrored) as predictors in the model and participant as a random intercept in the random effects term. We found a significant effect of Hemisphere (F(1, 1383.19) = 10.18, p < .001), a significant effect of Cluster (F(2, 1383.10) = 449.48, p < 0.001), and a significant effect of Navigation condition (F(3, 1384.94) = 7.85, p < 0.001) (Figure 5). We found a significant Cluster x Hemisphere interaction (F(2, 1383.18) = 4.49, p < 0.05) and a significant Cluster x Navigation condition interaction (F(6, 1383.12) = 5.53, p < 0.001).

**Figure 5.**
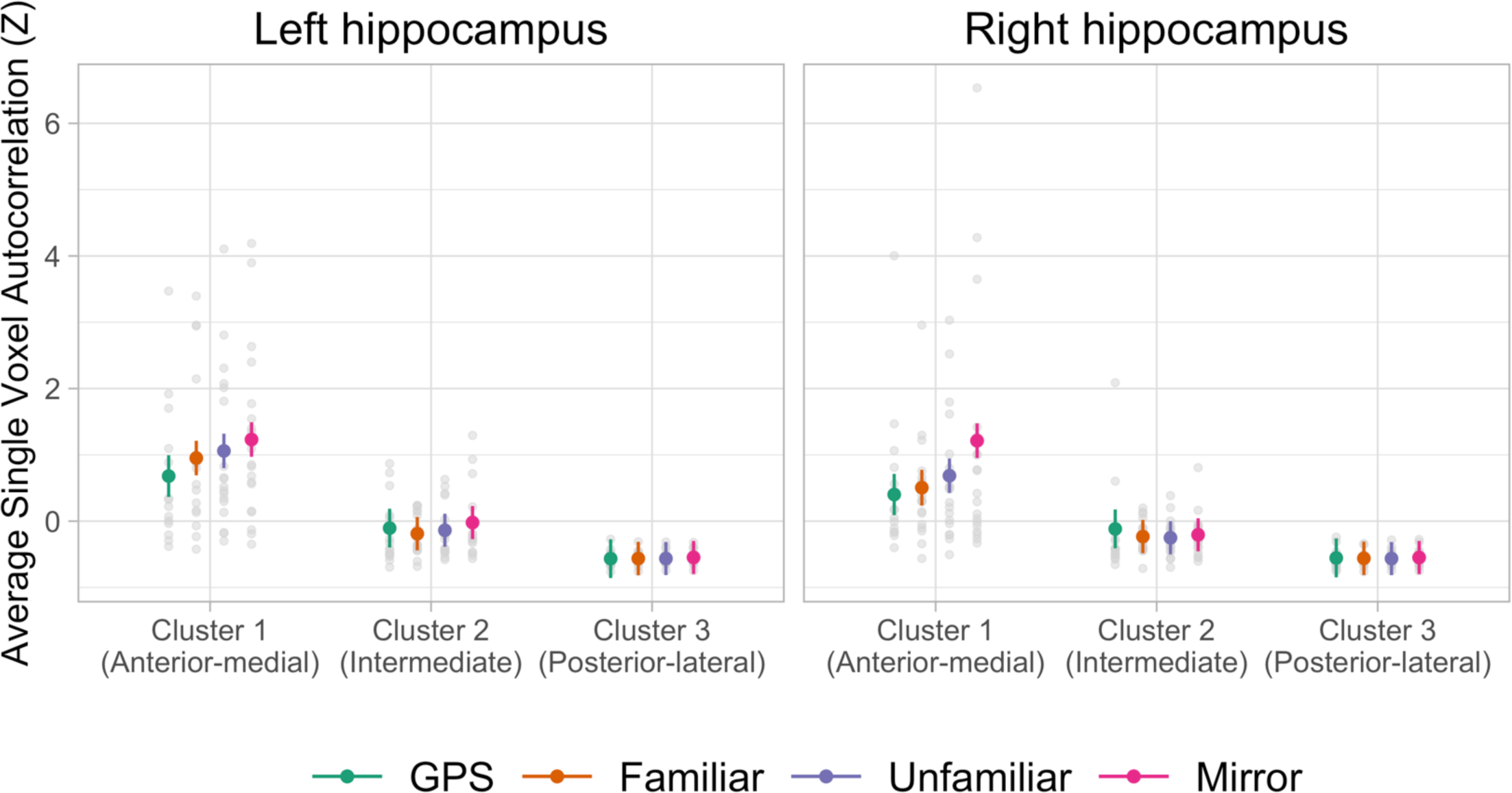
Effect of navigational condition on average single voxel autocorrelation. Average single voxel autocorrelation (lag 1) per cluster for each navigation condition. Across both hemispheres, Cluster 1 had greater average single voxel autocorrelation compared to Cluster 2 and 3. The average single voxel autocorrelation in Cluster 1 was modulated by navigational condition (in both left and right hemispheres). The autocorrelation of Cluster 1 was greatest when participants navigated Mirrored routes compared to GPS Familiar, and Unfamiliar routes. The autocorrelation for Unfamiliar routes was greater than GPS routes. There was no significant difference between Unfamiliar and Familiar, however the Unfamiliar routes were numerically greater than Familiar (in both hemispheres). There was no significant difference between the two easiest routes (Familiar and GPS). Error bars denote standard error.

A post hoc analysis of the main effect of Hemisphere revealed that the single voxel autocorrelation was greater in the left hippocampus compared to the right hippocampus (t(1383) = 3.19, p < 0.01). Pairwise comparisons of the clusters suggest that autocorrelation was greatest in Cluster 1 compared to Cluster 2 (t(1383) = 21.01, p < 0.001) and Cluster 3 (t(1383) = 29.45, p < 0.001).

Cluster 2 was greater than Cluster 3 (t(1383) = 8.96, p < 0.001). Analysis of the main effect of Navigation condition (collapsed across Hemisphere and Cluster) revealed that autocorrelation was significantly greater for the Mirrored compared to GPS (t(1387) = 3.86, p < 0.001), compared to Familiar (t(1383) = 4.14, p < 0.001) and compared to Unfamiliar (t(1383) = 3.11, p < 0.05). There was no significant difference between Familiar and Unfamiliar conditions and no significant difference between Familiar and GPS.

An analysis of the Cluster x Hemisphere interaction revealed that differences between hemispheres was driven by differences in Cluster 1, where the left hemisphere was greater than the right (t(1383) = 3.93, p < 0.001)). There were no significant differences between Cluster 2 and Cluster 3 in the left and right hemispheres. Further, the Navigation condition x Cluster interaction revealed that there were significant differences between navigation conditions, particularly in Cluster 1. There were no significant differences between navigation conditions in Cluster 2 or Cluster 3. Therefore, we report the pairwise comparisons for the different navigation conditions in Cluster 1, collapsed across hemispheres.

In Cluster 1, the routes rated as most difficult, the Mirrored routes, had higher autocorrelation compared to GPS (t(1384) = 6.22, p < .001), Familiar (t(1383) = 5.45, p < .001), and Unfamiliar (t(1383) = 3.95, p < .001). The Unfamiliar routes had higher autocorrelation compared to GPS (t(1384) = 3.03, p < .001). There was no significant difference between the two easier routes (Familiar and GPS) (p = 0.33). There were no significant differences in autocorrelation between Unfamiliar and Familiar (p = 0.39), however Unfamiliar routes were numerically greater than Familiar (in both hemispheres). This finding suggests that more difficult navigation conditions were related to increases in the autocorrelation in the anterior-medial hippocampus (Cluster 1).

#### Temporal SNR Control Analysis

We ran a linear mixed effects model on the average autocorrelation per cluster (lag 1) with Hemisphere (left, right), Cluster(1,2,3), Navigation condition (GPS, Familiar, Unfamiliar, Mirrored), and temporal SNR per cluster as fixed effects predictors. We found a significant effect of Cluster (F(2,1370.90) = 99.53, p < 0.001), a significant effect of Navigation condition (F(3,1359.73) = 4.74, p < 0.01), and a significant effect of temporal SNR (F(1,1376.58) = 840.80, p < 0.001). Importantly we found an interaction of Cluster and Navigation condition (F(6,1359.73) = 2.55, p < 0.05). This suggests that Cluster and Navigation condition significantly explained the variance in autocorrelation beyond temporal SNR.

Additionally, we ran two different model comparisons. First, we compared two models with Hemisphere, Navigation condition, and temporal SNR as predictors. One model included Cluster as a predictor and one model did not. We compared the models with a Chi square test and found that the model including Cluster was significant X2 (8, N = 19) = 689.47, p < 0.001, indicating that the model including Cluster was a better fit of the data than the model without. Next, we compared two models with Hemisphere, Cluster, and temporal SNR as predictors. One model included Navigation condition as a predictor and one model did not. We compared the models with a Chi square test and found that the model including Navigation condition was significant X^2^ (36, N = 19) = 140.827, p < 0.001, indicating that the model including Navigation condition was a better fit of the data than the model without. These results suggest that temporal SNR alone cannot explain the relationship between autocorrelation and navigation condition.

## Discussion

Here we present a novel autocorrelation measure to investigate intra- hippocampal and intra-entorhinal processing. We provide evidence of a medial-lateral gradient of autocorrelation in the hippocampus, as well as a posterior-medial and anterior-lateral gradient in the ERC. We found that voxels in the anterior-medial hippocampus have a highly correlated, slower changing signal, whereas voxels in the posterior-lateral hippocampus have a less correlated, faster changing signal (Figure 2) (Brunec, Bellana, et al., 2018; Raut et al., 2020). Our study highlights the importance of examining the medial-lateral axis of the hippocampus, which has previously been an under-studied feature of hippocampal organization. We find novel evidence for a continuous gradient in the ERC, with greater autocorrelation in the posteromedial ERC and lower autocorrelation in the anterolateral ERC (Figure 3). Lastly, the present study is the first to show that gradients of single voxel autocorrelation in the hippocampus are related to behavior during navigation. Specifically, autocorrelation in the anterior-medial hippocampus increased for difficult routes and was the greatest in the left hemisphere (Figure 5), whereas autocorrelation in the posterior-lateral hippocampus was similar across routes. These findings support the hypothesis that autocorrelation gradients are driven by intrinsic functional properties of the hippocampus and entorhinal cortex and suggest that these gradient patterns are not being driven by reciprocal connectivity or physiological noise (Figure 1D).

Our data-driven approach — which allows voxels to cluster according to their single voxel autocorrelation, uncovered a multidimensional gradient in both the anterior-posterior and medial-lateral axes in both the hippocampus and ERC (Figure 2 & 4). In the hippocampus, the anterior-posterior axis has been studied with respect to its role in representing graded information, for example coarse-grained to fine-grained information (Poppenk et al. 2013; Strange et al., 2014), large to small spatial distances (Evensmoen et al., 2013; Nielson et al., 2015; Peer et al., 2019;) and long to short temporal distance (Bellmund et al., 2019; Nielson et al., 2015). Investigations of representational differences along the medial-lateral axis, however, have been limited because prior work has used predefined anatomical segmentations limited to the anterior and posterior portions of the long axis of the hippocampus. Our single voxel autocorrelation method is not restricted by predefined ROIs and proves to be a more precise measure that detects subtle differences in signal along the medial-lateral axis that have been previously overlooked and that are modulated by navigational difficulty. Moreover, our findings are consistent with recent evidence of an anterior-medial hippocampal cluster that has been identified in resting-state MRI studies (Dalton et al., 2021, Thorp et al. 2021) and functional fMRI studies, which find that the anterior-medial hippocampus plays a distinct role in a range of cognitive and episodic memory tasks (Addis et al., 2012; Dalton et al., 2018; Lee et al., 2013; Zeidman et al., 2015, 2016). In addition to the hippocampus, we found similar distinctions in the ERC. We observed a gradient of single voxel autocorrelation organization, such that greater single voxel autocorrelation was observed in the posterior-medial region and lower single voxel autocorrelation in the anterolateral region of the ERC (Figure 3). This gradient is consistent with previous neuroimaging investigations of ERC which used high-resolution fMRI and functional connectivity to define distinct subregions within the human ERC (Maass et al., 2015; Navarro Schröder et al., 2015). Our analytic technique, however, goes beyond this prior work by demonstrating, for the first time, continuous gradients of autocorrelation in the ERC.

The present study demonstrates that the autocorrelation of the fMRI signal is not just global noise, but instead carries meaningful information about brain function that is directly related to behavior. Autocorrelation is frequently characterized as noise that masks meaningful signals and is unrelated to neuronal activity and cognition, therefore we included steps in our analyses to control for noise and artifacts that might be driving the autocorrelation signal (see Materials and Methods for details). A well-known source of autocorrelation of the fMRI signal comes from convolving the signal with the HRF (Friston et al., 1995). Recent research found that, while the autocorrelation signal mostly originates from the HRF, neural activity also contributes to the autocorrelation signal (Arbabshirani et al., 2019). Moreover, autocorrelation distinguished between global differences in cognitive state (task vs. rest) and mental state (healthy vs. schizophrenia) and these differences could not be accounted for by the HRF, physiological noise or motion (Arbabshirani et al., 2019). We therefore believe that the autocorrelation of the HRF alone cannot account for the findings presented here and that the autocorrelation is a measure of meaningful neural activity.

The interpretation of single voxel autocorrelation as a behaviorally relevant signal is corroborated by studies that use different measures of within-subject moment-to-moment signal and then relate these measures to cognitive performance. Specifically, research measuring changes in signal variability, as assessed by the standard deviation in BOLD signal over time, found that variability was related to task performance (see Garrett et al., 2013 for a review). For example, in younger adults, standard deviation of the BOLD signal was modulated by cognitive task demand, and as task difficulty increased, the variability decreased across the brain (Garrett, McIntosh, and Grady, 2014). This finding is consistent with our findings from Dataset2, wherein increased autocorrelation (i.e., lower variability) was related to increases in navigational difficulty. Our technique goes beyond prior methods by measuring the stability of a single voxel’s signal across multiple successive timepoints. This allows us to examine how the signal of individual voxels is maintained over an extended window of time and shows that this measure is directly related to behavior.

Recent research suggests that autocorrelation might be a global organizing principle and reflects intrinsic functional hierarchies in the brain (Irish & Vatansever, 2020; Raut et al., 2020). For example, an analysis of resting state fMRI data calculated the autocorrelation decay in single voxels across a temporal window (0-8 seconds) and found a significant large-to-small timescale gradient along the anterior-posterior axis in the hippocampus (Raut et al., 2020), which is consistent with reports by Brunec, Bellana, et al. (2018). While current studies of autocorrelation cannot address the direct link between the autocorrelation gradients and behavior, our work suggests that autocorrelation can be used to discriminate between cognitive states that are uniform across the brain, leaving open the question of how autocorrelation gradients in specific brain regions might be related to differences in cognition during a behavioral task. Our analysis technique demonstrated novel gradients during resting state and can also be applied to task related activation to reveal their relation to on-going behavior and is the first to show that changes in single voxel autocorrelation gradients are directly related to changes in difficulty during a navigation task.

Anterior hippocampal voxels are more stable across time compared to the posterior hippocampus, which might enable the anterior hippocampus to maintain prior information across time during goal-directed navigation (Brunec, Bellana, et al., 2018). Our method proved to be a more sensitive measure than previous techniques (e.g., Brunec, Bellana et al., 2018) because we were able to show differences in autocorrelation across navigation conditions. More specifically we found that the autocorrelation in the anterior-medial hippocampus increased during navigation of difficult routes (Figure 5). The autocorrelation signal may reflect the mechanism by which the hippocampus holds onto the past and carries it forward during navigation when we are in unfamiliar or unpredictable environments. For example, when navigating an unfamiliar route to a distant goal, the local details of the environment might not be helpful to orient oneself in relation to the goal; it may be more efficient, therefore, to keep in mind a coarser, overall map of the environment with information about steps already taken in order to reach the goal destination successfully. This large-scale representation may not be as useful to keep online during navigation of well-known or familiar routes where local details are sufficient for orienting and navigating to the goal, which could explain the decreased single voxel autocorrelation in the signal throughout the familiar routes (Figure 5). This hypothesis is supported by previous research which has shown that the anterior hippocampus plays an important role in representing larger spatial and temporal distances (Evensmoen et al., 2013; Nielson et al., 2015) as well as representing coarser-grained, global representations (Collin et al., 2015).

We found that single voxel autocorrelation in the anterior-medial hippocampal cluster was higher in the left hemisphere compared to the right hemisphere (Figure 5). It is still unclear whether this is representative of a stable difference in autocorrelation between the hemispheres or whether this reflects different types of information that are engaged across the two hemispheres during navigation. Future research is needed to determine the nature of this hemispheric difference.

Another non-mutually exclusive possibility is that the single voxel autocorrelation is representative of predictions that are cast into the future. The notion that increased temporal similarity is indicative of an extended spatiotemporal representation is supported by recent work investigating the predictive horizons along the hippocampal anteroposterior axis during navigation (Brunec & Momennejad, 2022). Brunec and Momennejad (2022) found that as participants virtually navigated familiar, real-world routes (a subset of the familiar routes presented here), hippocampal activity was related to a hierarchical scale of horizon representations, in which the posterior hippocampus represented steps closer in the future trajectory (∼25m) while the anterior hippocampus represented steps further in the future trajectory (∼175m). It is possible, therefore, that the single voxel autocorrelation we observed helps represent an upcoming navigational trajectory, with immediate goals represented in posterior-lateral regions and more distal goals in the anterior-medial hippocampus. Our method and findings open the door for future studies using high resolution neuroimaging in combination with a task that parametrically modulates the amount of information that is carried over time in both predictable (familiar) and unpredictable (unfamiliar) environments to uncover content that is carried forward via the autocorrelation signal.

Although we were not able to relate the autocorrelation in ERC to behavior due to the resolution of the navigation data, if we apply the same logic we used for the hippocampus, our findings are consistent with the notion that alERC codes for local details and perceptual aspects of experience, whereas the pmERC codes for global contexts. Specifically, the (antero-)lateral ERC has been linked to fine-grained temporal processing (Montchal et al., 2019; Tsao et al., 2018) and to processing of object-context and within-object details (Yeung et al., 2017; 2019). The low autocorrelation we observed in the alERC might indicate faster updating of moment-to-moment changes and therefore support fine-grained representations. Future investigations can use our method to analyze continuous changes along both anterior-posterior and medial-lateral axes of the ERC without being restricted to anatomical subfield segmentations, perhaps revealing a more nuanced understanding of the organization of the ERC. We observed two consistent clusters in the left hemisphere and three consistent clusters in the right hemisphere, which suggests that in this dataset there was more variability in the left ERC intermediate cluster. Future research is needed to determine whether this is a stable property of the temporal organization of the left ERC that can be replicated across other datasets.

It is currently unclear how the posterior-medial and anterior-lateral subregions of the ERC are functionally related to the anterior and posterior regions of the hippocampus. In the present study we found that clusters in the anterior-medial hippocampus and posteromedial ERC had high single voxel autocorrelation, whereas clusters in the posterior-lateral hippocampus and anterolateral ERC had low single voxel autocorrelation. These distinctions along the anterior-posterior and medial-lateral axes of the ERC are consistent with previous functional connectivity findings (Navarro Schröder et al., 2015), however functional connectivity and neuroanatomical studies in humans have been limited and do not find any clear differences between the anterior and posterior portions of the hippocampus with respect to their connectivity to different subregions in the ERC (Maass et al., 2015; Navarro Schröder et al., 2015). Functional connections between these regions might be evident in the scale of information processing in the hippocampus and ERC. For example, it is possible that the pattern of low single voxel autocorrelation in anterolateral ERC and posterior-lateral hippocampus supports fine-grained processing — precise temporal processing in the anterolateral ERC (Bellmund et al., 2019; Montchal et al., 2019) and local spatial details in the posterior hippocampus (Doeller et al., 2008; Evensmoen et al., 2013; Hirshhorn et al. 2012; Lee et al., 2012).

There are currently no clear neuroanatomical links between the anterolateral ERC and posterior-lateral hippocampus or the anterior-medial hippocampus and posteromedial ERC. There are, however, probable connections between the anterior ERC and lateral hippocampus and posterior ERC with medial hippocampus (Strange et al., 2014; Witter & Amaral 2020; Nilssen et al., 2019; Witter et al., 2017). Our results, therefore, open the door for future investigations to characterize more fully the nature of anterior and posterior hippocampal signal dynamics in relation to the entorhinal subregions in humans and in relation to other structures, such as prefrontal cortex (Barredo et al., 2015; Vaidya & Badre, 2020).

The results presented here reveal two continuous gradients along the anterior-posterior and medial-lateral axes in the hippocampus and ERC. One outstanding question is whether there is new information that can be gained by investigating the two autocorrelation gradients separately, or whether the information they represent is redundant. Another outstanding question is whether our novel single voxel autocorrelation method can be applied with shorter timescales so that they can be used with event-related designs. Here we use the entire timecourse of the voxel’s activity to calculate the single voxel autocorrelation throughout the entire run, but it remains to be seen whether we can adapt our method to examine how autocorrelation changes over shorter time windows. This would allow us to ask new questions about what kind of information is being carried in the autocorrelation signal during discrete or shorter events and at event boundaries, which are known to trigger changes in hippocampal activity associated with integration of information across events (Dubrow & Davachi, 2013; Ezzyat & Davachi, 2014). Finally, this method can be used to investigate differences in autocorrelation within subfields of the hippocampus. For example, it has been proposed that CA1 is implicated in integrating information in memory, whereas DG/CA3 which mediates pattern separation may be more implicated in making fine distinctions in memory (Kyle et al., 2015; Leutgeb et al., 2004; Schapiro et al., 2017; Yassa & Stark, 2011).

Integration processes in CA1, therefore, might be supported by voxels with high single voxel autocorrelation while separation processes in DG/CA3 might be better supported by low single voxel autocorrelation. Moreover, recent evidence also suggests that resting-state functional connectivity gradients in the hippocampus along the anterior-posterior and medial-lateral axes are closely associated with the microstructure of hippocampal subfields (deWael et al., 2018). Future research using our method and high-resolution fMRI is needed to test these differences within subfields.

Our studies were inspired initially by single-unit recording studies in rodents (Brun et al., 2008; Cavanagh et al., 2016; Gothard et al.,1996; Kjelstrup et al., 2008; Maurer et al., 2005). We believe our findings, however, have gone beyond replicating the rodent findings in humans, a worthy task in its own right, but extended the findings to the point that they can now be used to inform future studies in rodents and humans. We provide some examples in which this is the case. For example, our method enabled us to find differences in autocorrelation along the anterior-posterior and medial-lateral axes in the entorhinal cortex, which have only been examined in a restricted region in rodents (Brun et al., 2008). Our findings are consistent with neuroanatomical and neurophysiological divisions in that structure (human: Maass et al., 2015; monkey: Witter & Amaral, 2021; rat: Witter et al., 2017). Further, our findings are consistent with recent evidence from resting-state fMRI of two functional connectivity gradients in the hippocampus: a primary gradient along the anterior-posterior axis and a secondary gradient along the medial-lateral axis (deWael et al., 2018). Second, although activity of a single voxel that is comprised of thousands of neurons, may be considered to be a coarser unit of analysis than recordings from single units, it may be the case that it is the operation of a population of these neurons that is most closely linked to organizational temporal dynamics. It is the gradients revealed by autocorrelation at the single voxel level that enabled us to link hippocampal dynamics to behavior. In addition, we were able to segment the populations into clusters, suggesting subdivisions that would not be evident at the single-unit level. It would be worthwhile to determine whether similar clusters are found in rodents and examine their functional significance. Similar analyses at the population-level in rodents may yield information about the relation of neural dynamics to higher-level memory representations and goals, an enterprise that is just beginning (Jacob & Josselyn, 2020; Morrissey et al., 2017).

Our results provide compelling evidence for a gradation of single voxel autocorrelation in the hippocampus and ERC. Our single voxel method proved to be a fine-grained measure that revealed subtleties in the spatial organization of autocorrelation, going beyond prior methods, and allowed us to observe graded signals along anterior-posterior and medial-lateral axes in both regions. Further, we show for the first time that differences in single voxel autocorrelation gradients in the hippocampus can be directly related to differences in difficulty during a virtual navigation task, thus opening the door for future research to ask new questions of the autocorrelation signal and uncover how it is related to behavior.

## Acknowledgments

We thank Menno Witter for his feedback on an earlier draft of this paper. Dataset 1 was provided by the Human Connectome Project, WU-Minn Consortium (Principal Investigators: David Van Essen and Kamil Ugurbil; 1U54MH091657) funded by the 16 NIH Institutes and Centers that support the NIH Blueprint for Neuroscience Research; and by the McDonnell Center for Systems Neuroscience at Washington University. We thank Jason Ozubko for his contributions to the experiment conceptualization of and data collection for Dataset 2. MDB was supported by the Natural Sciences and Engineering Research Council Discovery Grant (RGPIN-2020-05747), a James S McDonnell Scholar Award, an Early Researcher Award from the Ontario Ministry of Development and Innovation, and a Canada Research Chair. MM was supported by the Canadian Institutes of Health Research Grant (MOP 125958).

