## Supplemental Materials for "Single voxel autocorrelation uncovers gradients of temporal dynamics in the hippocampus and entorhinal cortex during rest and navigation"

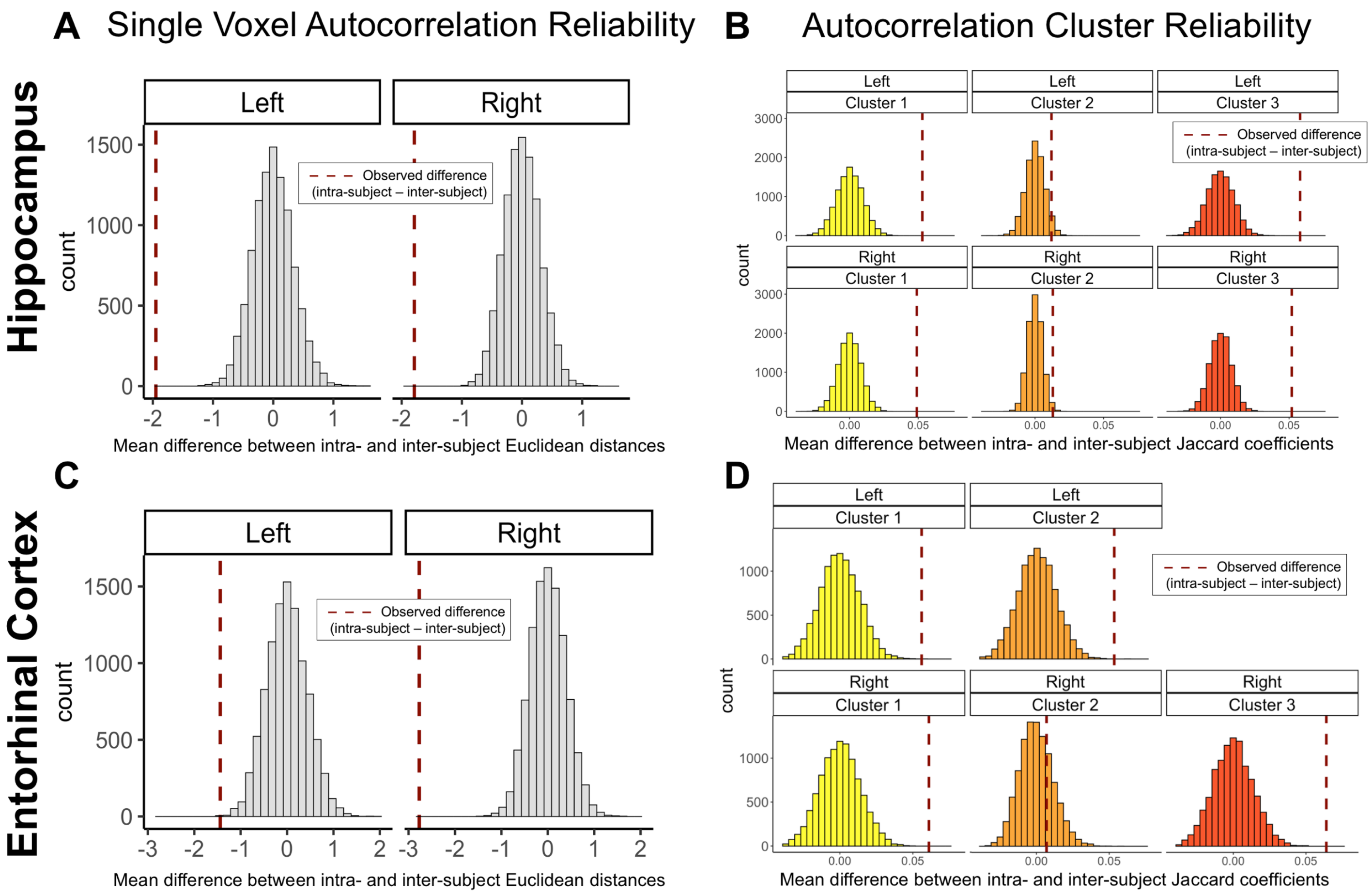


**S1. Hippocampal and Entorhinal cortex reliability measures. A,C) Single Voxel Autocorrelation Reliability.** Distribution of shuffled and permuted mean difference of intra- and inter-subject Euclidean distances for the (A) hippocampus and (C) entorhinal cortex. Dashed lines represent the observed mean difference between intra- and inter-subject Euclidean distance. Significant negative values indicate that single voxel autocorrelation values were more similar within an individual than across individuals. **B, D) Autocorrelation Cluster Reliability.** Distribution of shuffled and permuted mean difference of intra- and inter-subject Jaccard coefficients for each cluster. Dashed lines represent the observed difference between intra- and inter-subject Jaccard coefficients for each cluster. (B) In both hemispheres of the hippocampus, Clusters 1 and 3 were more reliable within individuals compared to Cluster 2. (D) In the entorhinal cortex, Cluster 1 and Cluster 2 were reliable within individuals in the left hemisphere, whereas Cluster 1 and 3 were reliable within individuals in the right hemisphere.
